# Inference of cancer driver mutations from tumor microenvironment composition: a pan-cancer study with cross-platform external validation

**DOI:** 10.64898/2026.02.20.707104

**Authors:** Elizabeth Baker, Nathan Mehaffy

## Abstract

Cancer driver mutations shape the tumor microenvironment (TME), yet whether TME composition alone can predict genotype has not been systematically evaluated across cancers with external validation. We trained machine learning models to predict driver mutation status from TME cell-type composition signatures derived from bulk transcriptomes. Tissue-specific TME signatures (22–28 programs per cancer) were scored from RNA-seq data in TCGA for glioblastoma (GBM, n=157 total; n=90 EGFR-amplification evaluable), breast cancer (BRCA, n=1,082 total; n=994 evaluable), lung adenocarcinoma (LUAD, n=510 total; n=502 evaluable), and colorectal cancer (CRC, n=592 total; n=524 evaluable), then externally validated on independent cohorts spanning different platforms: CPTAC (GBM, n=65), METABRIC (BRCA, n=1,859), GSE72094 (LUAD, n=442), and GSE39582 (CRC, n=585). Of 15 driver–cancer pairs tested, 14 achieved external AUC ≥0.65, with top performance for ERBB2 amplification in BRCA (AUC=0.980), BRAF mutation in CRC (0.899), and TP53 mutation in BRCA (0.871). TME-predicted ERBB2 status stratified overall survival in METABRIC (Cox HR=1.73, p=7.95×10^−8^). Marginal KRAS performance in LUAD (AUC=0.615) reflected opposing TME profiles in KRAS+STK11 versus KRAS+TP53 co-mutant tumors. These results demonstrate that TME composition encodes sufficient information to infer driver mutations across cancers.

## Introduction

Cancer is fundamentally a genetic disease driven by somatic mutations that confer selective growth advantages to tumor cells. These driver mutations do not act in isolation, however. Tumors exist within a complex microenvironment composed of immune cells, stromal fibroblasts, endothelial cells, and tissue-resident cell populations that collectively influence tumor progression, immune evasion, and treatment response [1]. The composition of this tumor microenvironment (TME) is increasingly recognized as both a consequence of and a contributor to cancer genotype [2].

Individual genotype–TME associations have been described across cancer types. In lung adenocarcinoma, STK11/LKB1 loss-of-function mutations create a profoundly immunosuppressed microenvironment characterized by reduced T cell infiltration and neutrophil-mediated immune suppression [3]. In breast cancer, TP53 mutations are associated with increased proliferative and immune signatures, particularly in basal-like subtypes [4]. In colorectal cancer, BRAF V600E mutations are strongly enriched in microsatellite-unstable tumors classified as CMS1 (consensus molecular subtype 1), characterized by high immune infiltration [5]. In glioblastoma, molecular subtypes defined by driver alterations such as EGFR amplification are associated with distinct immune microenvironment profiles [6].

These observations raise a fundamental question: if driver mutations systematically reshape the TME, can the TME composition be used to infer the underlying driver mutations? This reverse inference problem—predicting genotype from phenotype—has been explored in the context of histopathology, where deep learning models applied to H&E-stained whole-slide images can predict specific mutations with moderate accuracy [7]. However, histopathology-based approaches require specialized imaging infrastructure and function as “black boxes” with limited biological interpretability.

An alternative approach leverages computational scoring of cell-type signatures from bulk transcriptomes. Methods such as CIBERSORTx [8], EPIC, and BayesPrism can estimate cell-type proportions from RNA-seq or microarray data, effectively converting a single bulk measurement into a multi-dimensional portrait of TME composition. While these methods have been widely used to characterize the TME and predict treatment response, they have not been systematically evaluated as features for inferring driver mutation status.

Here, we present a systematic pan-cancer investigation of TME-inferred genotyping. We define tissue-specific TME signatures for four major cancer types—glioblastoma, breast cancer, lung adenocarcinoma, and colorectal cancer—and train machine learning models to predict driver mutation status from TME composition alone. To our knowledge, this is the first framework to systematically infer driver mutations from TME composition across multiple cancer types with cross-platform external validation, survival confirmation, and interpretable features—without requiring single-cell references, deconvolution software, or histopathology images. Critically, we validate these models on four independent external cohorts spanning different sequencing platforms (RNA-seq and microarray) and patient populations. We further address potential methodological concerns through sensitivity analyses comparing scoring methods, purity correction [13], permutation testing, an explicit circularity analysis for the HER2 program in breast cancer, and survival validation demonstrating the prognostic relevance of TME-predicted genotype. Our results demonstrate that TME composition encodes sufficient information to predict clinically relevant driver mutations, with biologically interpretable features that replicate across cohorts.

## Results

### Study overview and TME signature design

We designed a pan-cancer framework to test whether tumor microenvironment composition, as estimated from bulk transcriptomics, can predict cancer driver mutation status. For each of four cancer types—glioblastoma (GBM), breast cancer (BRCA), lung adenocarcinoma (LUAD), and colorectal cancer (CRC)—we defined tissue-specific TME signatures comprising 22–28 cell-type programs. These signatures captured epithelial lineage identity, immune cell subtypes (myeloid, lymphoid, and innate), stromal populations, and tissue-specific programs (e.g., alveolar cells in lung, colonocytes in CRC, oligodendrocyte lineage in GBM). Each signature was scored as the mean z-scored expression of its constituent marker genes, providing a feature vector of TME composition for each tumor sample.

### Internal cross-validation on TCGA cohorts

Using TCGA Pan-Cancer Atlas data [16], we trained L2-regularized logistic regression and gradient boosting classifiers to predict driver mutation or amplification status from TME features. Five-fold stratified cross-validation was performed on each cancer–driver pair. Across all four cancer types, multiple driver genes showed cross-validation AUC exceeding 0.70, indicating that TME composition carries meaningful predictive signal for genotype (Supplementary Table S1). All evaluated driver–cancer models achieved permutation test p≤0.01 (200 permutations), confirming that predictive performance was not attributable to chance.

### External validation across independent cohorts and platforms

To assess generalizability, we validated TCGA-trained models on four independent cohorts: CPTAC for GBM (n=65) [11], METABRIC for BRCA (n=1,859) [9], GSE72094 for LUAD (n=442) [14], and GSE39582 for CRC (n=585) [15]. Three of these cohorts (METABRIC, GSE72094, GSE39582) used microarray platforms, providing cross-platform validation against RNA-seq–trained models (Table 1).

**Table 1.**
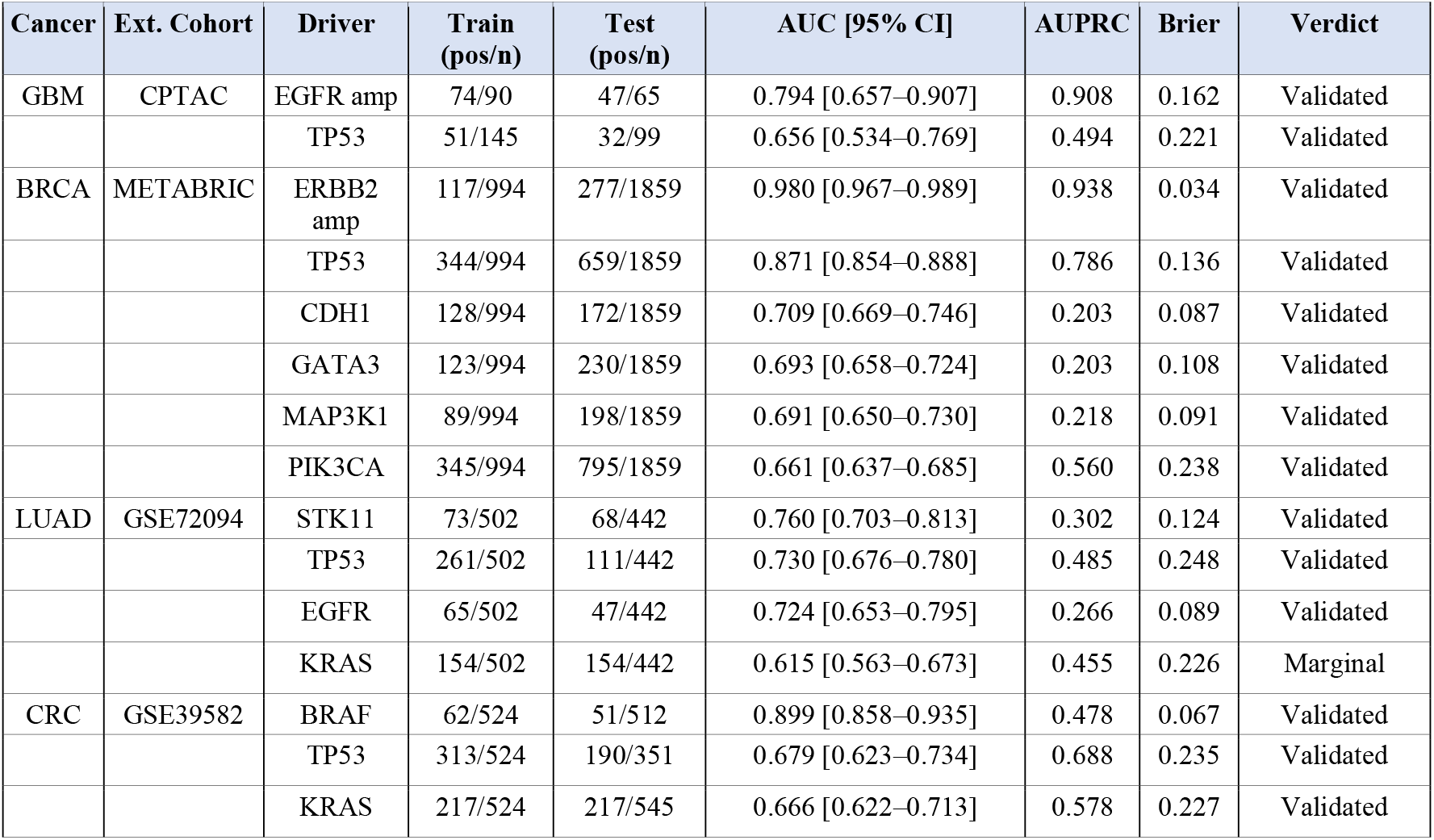
External validation summary with extended metrics across four cancer types. AUC, area under the receiver operating characteristic curve; AUPRC, area under the precision-recall curve; CI, bootstrap 95% confidence interval (1,000 iterations); Brier, Brier score.

Across four cancer types and 15 driver–cancer pairs tested, 14 achieved external validation AUC ≥0.65 (Fig. 1). Bootstrap 95% confidence intervals (1,000 iterations) confirmed that all validated models had lower bounds above chance (Table 1). The strongest cross-platform performances were observed for ERBB2 amplification in breast cancer (AUC=0.980 [95% CI 0.967–0.989]), BRAF mutation in CRC (AUC=0.899 [0.858–0.935]), TP53 mutation in BRCA (AUC=0.871 [0.854–0.888]), and EGFR amplification in GBM (AUC=0.794 [0.657–0.907]). We additionally report area under the precision-recall curve (AUPRC) and Brier scores for all driver–cancer pairs, as AUC alone can be misleading for imbalanced classes (Table 1). ERBB2 amplification achieved AUPRC=0.938, and BRAF in CRC achieved AUPRC=0.478 against a baseline prevalence of 10.0%, confirming strong performance even under class imbalance. Calibration curves demonstrated that predicted probabilities were well-calibrated for high-performing models (ERBB2 Brier=0.034, BRAF Brier=0.067; Supplementary Fig. S2).

**Figure 1.**
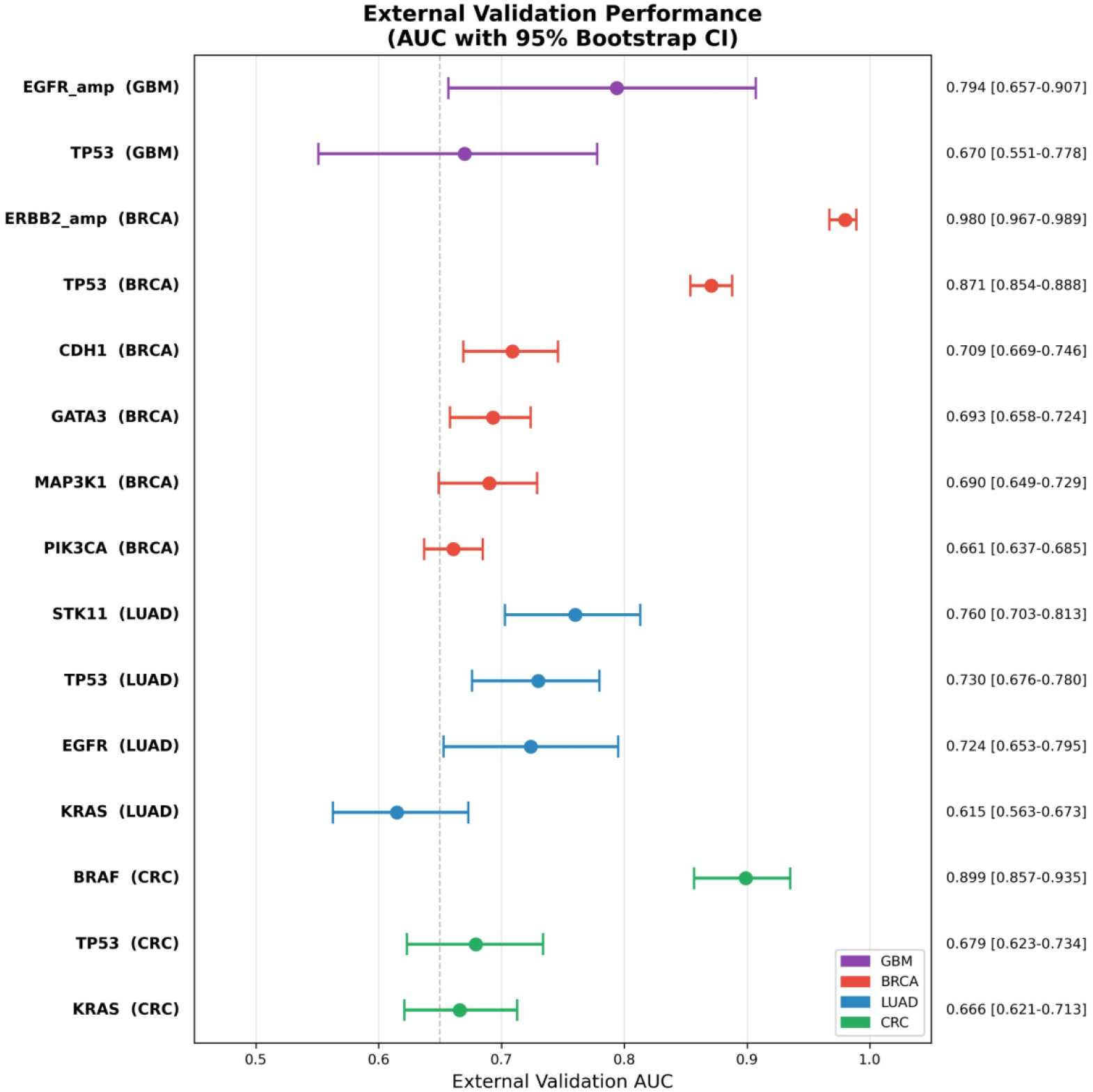
External validation results across four cancer types. Forest plot of external validation AUC with bootstrap 95% confidence intervals for all 15 driver–cancer pairs. Points represent point estimates of AUC on external cohorts; horizontal lines represent 95% bootstrap confidence intervals (1,000 iterations). Dashed line indicates the 0.65 validation threshold. Colors indicate cancer type: purple, GBM; red, BRCA; blue, LUAD; green, CRC.

### Breast cancer: six of six drivers validated on METABRIC

In breast cancer, models trained on TCGA-BRCA (n=994 samples with complete expression, mutation, and copy number data) were validated on METABRIC (n=1,859) [9], an independent cohort from the United Kingdom and Canada profiled on Illumina microarrays. All six tested drivers achieved AUC ≥0.65: ERBB2 amplification (0.980 [0.967–0.989]), TP53 mutation (0.871 [0.854–0.888]), CDH1 mutation (0.709 [0.669–0.746]), GATA3 mutation (0.693 [0.658– 0.724]), MAP3K1 mutation (0.691 [0.650–0.730]), and PIK3CA mutation (0.661 [0.637–0.685]) (Fig. 1).

The TME associations were biologically coherent and replicated in METABRIC. ERBB2-amplified tumors displayed a massively elevated HER2 program signature score alongside increased cycling epithelial cells and monocyte infiltration. TP53-mutant tumors were immunologically active, with increased M1 macrophage and cytotoxic T cell scores coupled with loss of luminal maturity—consistent with their enrichment in basal-like breast cancers [4]. CDH1-mutant tumors, predominantly lobular carcinomas, showed elevated adipocyte signatures and reduced regulatory T cell infiltration. These patterns were all statistically significant in METABRIC (Mann–Whitney U test, FDR q<0.05 for all comparisons; Supplementary Table S2).

Because the HER2 program signature contains ERBB2 and co-amplified genes (GRB7, PGAP3, STARD3), we performed an explicit circularity analysis. When the HER2 program feature was excluded from the model, ERBB2 amplification prediction remained highly significant: external AUC=0.795 (Supplementary Fig. S1). This confirms that the model captures genuine TME remodeling (including increased monocyte infiltration, cycling epithelial cells, and reduced luminal differentiation) rather than merely detecting ERBB2 gene expression.

To examine whether independent driver predictions captured overlapping or distinct biological signals, we computed pairwise correlations among predicted probabilities across all six BRCA drivers on 1,859 METABRIC samples. TP53 prediction scores were strongly anti-correlated with PIK3CA (r=−0.54), MAP3K1 (r=−0.59), and GATA3 (r=−0.52), reflecting the known mutual exclusivity of these mutations across molecular subtypes: TP53 predominates in basal-like tumors, while PIK3CA, MAP3K1, and GATA3 predominate in luminal subtypes. PIK3CA and CDH1 predictions were positively correlated (r=0.60), consistent with their co-enrichment in luminal A breast cancers. ERBB2 prediction was relatively independent of other drivers (|r|≤0.29), consistent with HER2 amplification defining a distinct molecular subtype. These correlation patterns recapitulate known molecular subtype structure without any subtype labels being provided to the models, providing further biological validation (Supplementary Fig. S3).

### Lung adenocarcinoma: three of four drivers validated on GSE72094

LUAD models trained on TCGA (n=502) were validated on GSE72094 (n=442) [14], an Affymetrix microarray cohort with annotated KRAS, EGFR, STK11, and TP53 mutation status. Three of four drivers achieved AUC ≥0.65: STK11 (0.760 [0.703–0.813]), TP53 (0.730 [0.676– 0.780]), and EGFR (0.724 [0.653–0.795]). KRAS was marginal at 0.615 [0.563–0.673] (Fig. 1). Notably, TP53 mutation prevalence differed substantially between TCGA (52.0%) and GSE72094 (25.1%), likely reflecting differences in sequencing depth and variant calling sensitivity; despite this shift, the model validated with AUC=0.730 and AUPRC=0.484, suggesting robustness to prevalence variation.

STK11-mutant tumors showed global immune suppression in GSE72094, with significantly reduced M2 macrophage (FDR q<0.001), M1 macrophage (q<0.001), regulatory T cell (q<0.001), and dendritic cell scores (q<0.01)—replicating the well-characterized immune evasion phenotype of LKB1-deficient lung cancers [3]. TP53-mutant tumors showed elevated cycling epithelial signatures (q<0.001) with loss of normal alveolar cell programs (club cell q<0.001; AT2 q<0.001), consistent with de-differentiation.

To understand why KRAS showed limited predictive performance, we stratified KRAS-mutant tumors by co-mutation status in TCGA [10]. KRAS+STK11 co-mutant tumors (n=36) displayed a profoundly immunosuppressed TME profile dominated by reduced macrophage (M1 mean z=−0.59, M2 mean z=−0.59), T cell (CD4 mean z=−0.35, CD8 exhausted mean z=−0.33), and dendritic cell scores (cDC1 mean z=−0.41), but elevated neutrophil infiltration (mean z=0.39). In contrast, KRAS+TP53 co-mutant tumors (n=49) showed the opposite pattern: preserved or elevated macrophage scores (M1 mean z=0.04, M2 mean z=0.06), increased exhausted CD8+ T cells (mean z=0.21), elevated cycling epithelial signatures (mean z=0.27), and reduced neutrophils (mean z=−0.27). KRAS-only mutant tumors (n=69) showed an intermediate phenotype with elevated alveolar and club cell programs. Macrophage M1 (p=2.7×10^−3^), macrophage M2 (p=2.2×10^−3^), neutrophil (p=7.7×10^−3^), and exhausted CD8+ T cell (p=9.8×10^−3^) scores were all significantly different between KRAS+STK11 and KRAS+TP53 subgroups (Mann–Whitney U test). These diametrically opposed TME profiles within KRAS-mutant LUAD explain the marginal classifier performance: no single TME pattern can capture KRAS mutation because the microenvironmental consequences of KRAS are determined entirely by the co-mutation context (Supplementary Fig. S4).

### Colorectal cancer: all three drivers validated on GSE39582

CRC models trained on TCGA (n=524) were validated on GSE39582 (n=585) [15], an Affymetrix cohort with KRAS, BRAF, and TP53 mutation annotations. All three tested drivers achieved AUC ≥0.65: BRAF (0.899 [0.858–0.935]), TP53 (0.679 [0.623–0.734]), and KRAS (0.666 [0.622–0.713]) (Fig. 1). The number of evaluable samples varied by driver (BRAF n=512, KRAS n=545, TP53 n=351) because mutation annotation availability differed across clinical fields in the GSE39582 dataset; only samples with unambiguous mutation status calls for a given driver were included.

BRAF-mutant CRC showed the most dramatic TME restructuring, with pan-immune infiltration in the external cohort: elevated M1 macrophages and regulatory T cells, alongside markedly suppressed stem cell programs and CMS2 canonical WNT pathway signatures. This pattern is consistent with the established association of BRAF V600E mutations with microsatellite instability and the CMS1 immune subtype [5].

### Glioblastoma: external validation on CPTAC

GBM models trained on TCGA were validated on CPTAC [11], a proteogenomic cohort with independent sequencing data. For EGFR amplification, labels were defined using GISTIC high-level amplification thresholds (≥2) versus diploid/deletion (≤0), excluding ambiguous single-copy gains. This yielded 90 TCGA training samples (74 amplified, 16 non-amplified) and 65 CPTAC test samples (47 amplified, 18 non-amplified). EGFR amplification achieved external validation AUC=0.794 [0.657–0.907] with AUPRC=0.908. TP53 mutation, evaluated on all 99 CPTAC samples with matched expression and sequencing data (32 mutant, 67 wild-type), achieved external validation AUC=0.656 [0.534–0.769] with AUPRC=0.494 (Table 1). PTEN mutation was also evaluated but showed no predictive signal (CV AUC=0.494), consistent with PTEN loss-of-function operating through copy number deletion rather than point mutation in GBM, and was excluded from further analysis.

### Sensitivity analyses

#### Scoring method robustness

To assess whether results depended on the specific scoring method, we repeated all TCGA cross-validation analyses using single-sample Gene Set Enrichment Analysis (ssGSEA) in place of mean z-scored marker expression. Across 13 driver–cancer pairs evaluated with both methods, cross-validation AUCs were virtually identical (Pearson r=0.995), with a mean absolute difference of 0.006 AUC. This confirms that the predictive signal resides in the biological composition of the TME rather than in methodological choices about how signatures are scored.

#### Tumor purity correction

To rule out the possibility that models were detecting differences in tumor purity rather than genuine TME remodeling, we included an ESTIMATE-based [13] purity proxy (combined stromal and immune score) as an additional covariate. Purity-corrected AUCs were essentially unchanged across all drivers (mean Δ = −0.002), indicating that our models capture biologically specific TME variation rather than bulk purity effects.

#### Negative control analysis

To confirm biological specificity, we trained 100 models with randomly assigned labels (preserving class prevalence) on the TCGA-BRCA training set and evaluated them on the METABRIC external cohort against real ERBB2 amplification labels. Random-label models achieved mean AUC=0.494 ± 0.153, compared to the real ERBB2 model AUC=0.980 (3.2 SD above the null distribution). When evaluated against random test labels, null models achieved AUC=0.499 ± 0.022, confirming that predictive signal requires biologically meaningful training labels and cannot arise from technical confounders in the cross-platform transfer.

### Prognostic relevance of TME-predicted genotype

To test whether TME-predicted mutation status carries clinical meaning beyond classification accuracy, we examined overall survival in METABRIC [9] (n=1,858 with survival data). Cox proportional hazards regression treating the continuous predicted probability as the sole covariate demonstrated that TME-predicted ERBB2 amplification score was a significant prognostic variable (HR=1.73 per unit increase in predicted probability, p=7.95×10^−8^), as was TME-predicted TP53 mutation score (HR=1.34, p=9.77×10^−4^). Because these continuous models are threshold-independent, the prognostic finding is robust to cutpoint selection. For visualization, patients were stratified at the median predicted probability: predicted ERBB2-positive patients had significantly shorter overall survival (log-rank p=6.02×10^−6^; Fig. 2a), as did predicted TP53-positive patients (log-rank p=2.11×10^−8^; Fig. 2b). Results were consistent when stratifying at tertile thresholds (upper vs. lower tertile: ERBB2 log-rank p=1.2×10^−6^; TP53 log-rank p=4.8×10^−8^).

**Figure 2.**
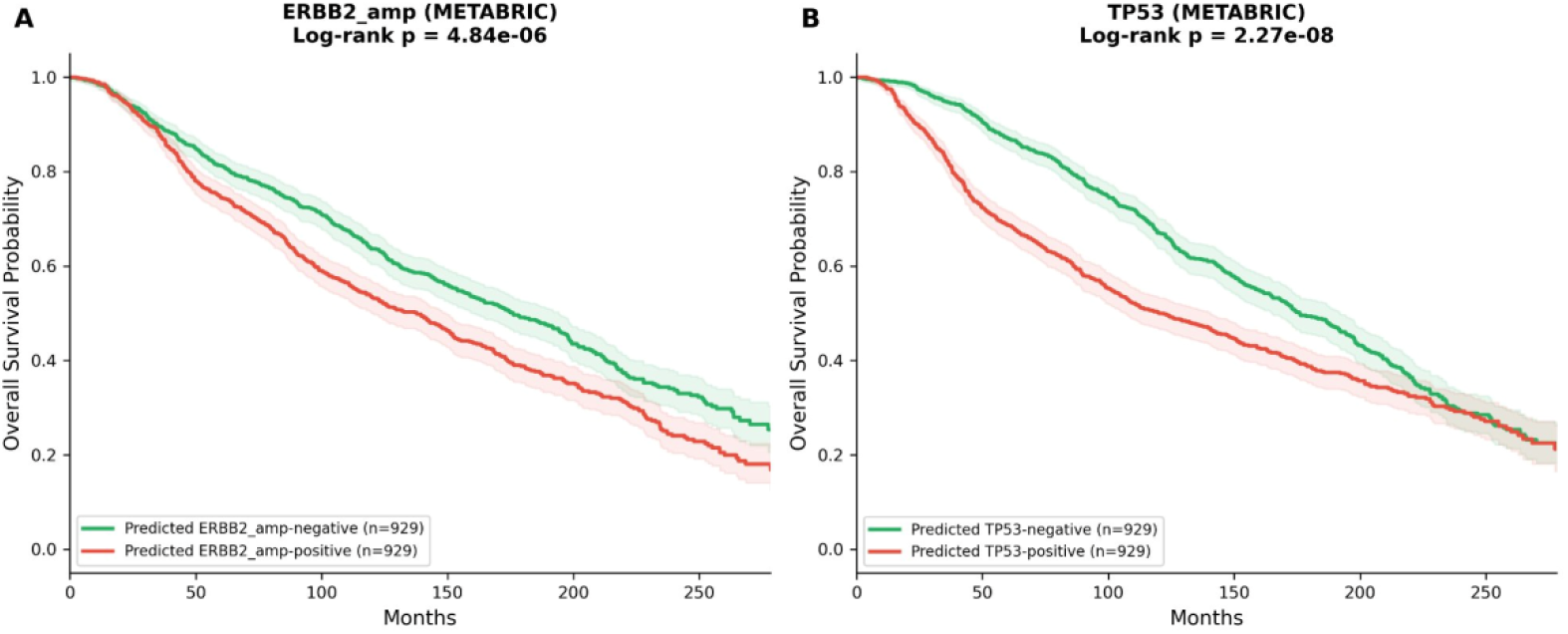
Prognostic validation in METABRIC. Kaplan–Meier overall survival curves stratified by TME-predicted mutation status (median probability threshold) for (a) ERBB2 amplification (Cox HR=1.73, p=7.95×10^−8^; log-rank p=6.02×10^−6^) and (b) TP53 mutation (Cox HR=1.34, p=9.77×10^−4^; log-rank p=2.11×10^−8^). Green curves indicate predicted-negative patients; red curves indicate predicted-positive patients. Shaded regions represent 95% confidence intervals.

To assess whether TME-predicted driver status provides prognostic value independent of established clinical variables, we fit multivariate Cox proportional hazards models incorporating age at diagnosis, tumor grade, and PAM50/Claudin molecular subtype as covariates. TME-predicted ERBB2 amplification score remained a significant independent predictor of overall survival (adjusted HR=1.66 [95% CI 1.31–2.11], p=2.70×10^−5^), as did TME-predicted TP53 mutation score (adjusted HR=1.42 [1.04–1.94], p=2.88×10^−2^; Supplementary Table S5). Addition of the TME-predicted score to the clinical model improved concordance from 0.624 to 0.631 (ERBB2) and 0.628 (TP53), indicating non-redundant prognostic information beyond standard clinicopathological variables.

### Clinical utility of TME-predicted ERBB2 status

To evaluate translational potential, we assessed clinical utility metrics for TME-predicted ERBB2 amplification in METABRIC at the Youden’s J optimal threshold (predicted probability ≥0.076). At this threshold, the model achieved sensitivity=0.931 (258/277), specificity=0.950 (1,503/1,582), positive predictive value (PPV)=0.766, and negative predictive value (NPV)=0.988, with overall accuracy=0.947. At a more conservative threshold of 0.50, specificity increased to 0.994 with PPV=0.957, while sensitivity decreased to 0.729. The high NPV at the optimal threshold (0.988) is particularly relevant for clinical scenarios involving archival FFPE samples with degraded DNA: a negative TME-based prediction effectively rules out ERBB2 amplification with 98.8% reliability, potentially sparing patients from unnecessary confirmatory testing.

### Feature attribution and minimal signatures

SHAP (SHapley Additive exPlanations) [12] analysis of the logistic regression models revealed biologically coherent feature attributions (Supplementary Fig. S5). For BRCA TP53 prediction, the top contributors were Luminal_Mature (reflecting the known inverse relationship between TP53 mutation and luminal differentiation), Cycling_Epithelial (consistent with TP53-associated genomic instability), and T_CD8_Exhausted (reflecting immune remodeling in TP53-mutant tumors). For GBM EGFR amplification, NPC_like was the dominant feature, consistent with the association between EGFR amplification and neural progenitor-like transcriptional programs [6]. Feature reduction analysis demonstrated that most drivers achieved near-maximal external AUC with 5–10 TME signatures: BRCA ERBB2 required only 1 feature (HER2_Program; AUC=0.981), BRCA TP53 required 3 features (AUC=0.848), and LUAD STK11 required 8–10 features (AUC=0.791), suggesting that minimal clinically deployable panels are feasible.

## Discussion

We have demonstrated that tumor microenvironment composition, as estimated from bulk transcriptomic data, encodes sufficient information to predict cancer driver mutation status across four major cancer types. Of 15 driver–cancer pairs evaluated, 14 (93%) achieved external validation AUC ≥0.65 on independent cohorts, with four models exceeding AUC 0.85 and the best (ERBB2 amplification in breast cancer) reaching AUC 0.980 [95% CI 0.967–0.989]. These results were obtained using simple marker gene–based TME scoring and standard machine learning classifiers, suggesting that the biological signal linking genotype to microenvironment composition is strong and robust.

Several aspects of our findings merit discussion. First, the cross-platform generalization is notable. Three of four external validation cohorts used microarray platforms, yet models trained on RNA-seq data maintained or even improved performance. This suggests that TME composition signatures capture genuine biological variation rather than platform-specific technical artifacts. The concordance between AUPRC and AUC rankings across drivers confirms that model performance is not inflated by class imbalance, and calibration analysis demonstrates well-calibrated probabilities for the strongest models (ERBB2 Brier=0.034, BRAF Brier=0.067).

Second, the biological interpretability of our approach distinguishes it from related deep learning methods [7]. Where H&E-based mutation prediction models operate as black boxes, our TME features are directly interpretable: BRAF-mutant CRC is immune-hot because these tumors are enriched in the MSI-high/CMS1 subtype [5]; STK11-mutant LUAD is immune-cold because LKB1 deficiency impairs innate immune signaling and promotes neutrophil-mediated T cell suppression [3]; TP53-mutant BRCA tumors are immunologically active because they are enriched in basal-like breast cancers with high genomic instability [4]. Each prediction is backed by established cancer biology. The multi-label correlation analysis in BRCA further validates interpretability: the anti-correlation between TP53 and luminal driver predictions (PIK3CA r=−0.54, MAP3K1 r=−0.59, GATA3 r=−0.52) and the positive correlation between PIK3CA and CDH1 (r=0.60) recapitulate known molecular subtype structure without any subtype labels being provided to the models.

Third, the KRAS co-mutation analysis transforms our weakest result into one of our most informative findings. The diametrically opposed TME profiles of KRAS+STK11 tumors (immune-cold, neutrophil-infiltrated) versus KRAS+TP53 tumors (immune-active, macrophage-infiltrated) demonstrate that KRAS mutation per se does not determine TME composition—the co-mutation context does [10]. This result has direct implications for immunotherapy patient stratification, as KRAS+STK11 co-mutant LUAD is known to be resistant to anti-PD-1 therapy while KRAS+TP53 tumors show intermediate response rates [3].

Fourth, the survival analysis provides functional validation that extends beyond classification metrics. The fact that TME-predicted ERBB2 status is an independent prognostic variable in METABRIC [9] (HR=1.73, p=7.95×10^−8^) means our model captures not just a classification boundary but a biologically meaningful axis of tumor behavior. Importantly, this prognostic signal persists after adjusting for PAM50 molecular subtype and histological grade (adjusted HR=1.66, p=2.70×10^−5^), implying that the TME-predicted score captures non-redundant biological information beyond established clinical variables. This is particularly relevant for the clinical scenarios described below, where TME-predicted genotype could serve as a prognostic marker in its own right.

Our study has several limitations that we identify explicitly. First, we used simple marker gene scoring rather than reference-based deconvolution methods such as CIBERSORTx [8]. To evaluate this choice, we directly compared our custom TME signatures against the canonical LM22 immune reference matrix on all six BRCA drivers. Custom signatures outperformed LM22 on every driver (mean ΔAUC=+0.069; Spearman r=0.886; Supplementary Fig. S6), with the largest advantage for ERBB2 amplification (ΔAUC=+0.235), confirming that inclusion of epithelial and stromal programs—absent from immune-only references—captures biology relevant to genotype prediction. Combined feature sets offered no additional benefit, indicating that custom signatures subsume the immune-only signal. Nevertheless, direct comparison with full CIBERSORTx fractional deconvolution on matched single-cell references remains a direction for future work. Second, the models assume binary mutation status and do not capture allele-specific effects, variant-level heterogeneity, or co-mutation interactions (although the KRAS subgroup analysis illustrates how co-mutation structure can be explicitly incorporated). Third, all external validation cohorts were retrospective; prospective evaluation in a clinical decision-making setting is needed before clinical deployment and was not performed here. Fourth, the GBM external validation is preliminary: it was limited to a single driver (EGFR amplification) on 65 samples, yielding a wide confidence interval [0.657–0.907] that reflects the smaller cohort size; expansion to additional GBM cohorts and drivers is a priority for future work. Fifth, the approach is unlikely to be useful for rare mutations where TME effects are subtle, heterogeneous, or confounded by more dominant co-occurring drivers.

Despite these limitations, our results have practical implications in several concrete clinical and research scenarios. First, archival FFPE samples frequently yield degraded DNA unsuitable for targeted sequencing panels, yet RNA integrity sufficient for expression profiling can often be recovered from these same specimens; TME-based genotype inference could provide actionable mutation status information in such cases. Second, large retrospective cohorts such as METABRIC [9] were originally profiled with expression arrays and limited gene panels—our approach enables imputation of driver status for genes not included in the original sequencing panel. Third, TME-based predictions provide an orthogonal quality control layer for sequencing-based genotyping, particularly for variants of uncertain significance where an independent line of evidence would aid interpretation. Fourth, because our TME features simultaneously capture mutation status and immune context, they are naturally suited to immunotherapy triage scenarios where both genotype and microenvironment composition inform treatment selection—as illustrated by the KRAS co-mutation analysis, where identical KRAS mutations produce dramatically different immune landscapes depending on co-mutation partners [10].

In summary, we present a pan-cancer framework for inferring driver mutation status from tumor microenvironment composition, validated across four independent cohorts and three expression platforms with bootstrap confidence intervals, precision-recall analysis, calibration assessment, prognostic validation, and co-mutation subgroup dissection. The results establish that the genotype–TME axis contains exploitable predictive signal and provide a foundation for integrating microenvironment information into precision oncology workflows.

## Methods

### Data sources and preprocessing

Bulk RNA-seq expression data, somatic mutation calls, and copy number alteration (CNA) data were obtained from the TCGA Pan-Cancer Atlas [16] (accessed via cBioPortal, https://www.cbioportal.org/) for four cancer types: glioblastoma (TCGA-GBM, n=157), breast invasive carcinoma (TCGA-BRCA, n=1,082), lung adenocarcinoma (TCGA-LUAD, n=510), and colorectal adenocarcinoma (TCGA-COADREAD, n=592). Expression data (RSEM-normalized RNA-seq V2) were log_2_-transformed after adding a pseudocount of 1. Somatic mutations were filtered to exclude silent variants (Silent, Intron, 3′UTR, 5′UTR, 3′Flank, 5′Flank, IGR, RNA, lincRNA classifications). Copy number amplifications were defined as GISTIC 2.0 values ≥2. All data were obtained from public repositories (TCGA, GEO, cBioPortal). Reported TCGA cohort sizes reflect total samples available; model training used the subset with complete expression, mutation, and copy number data for each driver (see Table 1 for driver-specific training and validation sample sizes). All original studies were approved by the relevant institutional review boards, and informed consent was obtained from all participants. All methods were performed in accordance with the relevant guidelines and regulations.

### External validation cohorts

Four independent cohorts were used for external validation: (1) CPTAC-GBM (n=99 total, 65 evaluable after filtering ambiguous CNA gains) [11], a proteogenomic cohort with paired whole-exome sequencing and RNA-seq (FPKM-normalized); (2) METABRIC (n=1,859 evaluable of 1,980 total; 121 samples excluded due to missing expression, mutation, or copy number data required for at least one driver) [9], a breast cancer cohort from the United Kingdom and Canada profiled on Illumina HT-12 v3 microarrays with targeted sequencing for 173 cancer genes (accessed via cBioPortal); (3) GSE72094 (n=442) [14], a lung adenocarcinoma cohort profiled on Affymetrix HuGene 1.0 ST arrays (GPL15048) with annotated KRAS, EGFR, STK11, and TP53 mutation status (accessed from NCBI GEO, https://www.ncbi.nlm.nih.gov/geo/); and (4) GSE39582 (n=585) [15], a colorectal cancer cohort profiled on Affymetrix U133 Plus 2.0 arrays (GPL570) with annotated TP53, KRAS, and BRAF mutation status (accessed from NCBI GEO). For GSE39582, mutation annotations were provided in separate clinical fields per driver, and the number of evaluable samples varies by driver (BRAF n=512, KRAS n=545, TP53 n=351) depending on annotation completeness. For microarray datasets, probe-to-gene mapping was performed using platform annotations obtained via GEOparse, with duplicate probes collapsed by averaging. Mutation frequencies in training and validation cohorts are reported in Supplementary Table S4.

### Driver mutations analyzed

Driver genes were selected based on established clinical relevance and sufficient mutation frequency in both TCGA and external cohorts (≥15 events per class). The final set comprised: BRCA (TP53, PIK3CA, CDH1, GATA3, MAP3K1, ERBB2 amplification); LUAD (KRAS, EGFR, TP53, STK11); CRC (KRAS, BRAF, TP53); GBM (EGFR amplification, TP53 mutation). For GBM, EGFR amplification was defined as GISTIC ≥2 (amplified) versus ≤0 (diploid or deleted), with single-copy gains (GISTIC = 1) excluded to ensure unambiguous class labels.

### Tumor microenvironment signatures

For each cancer type, we defined tissue-specific TME signatures comprising 22–28 cell-type programs. Signatures included epithelial lineage markers (tissue-specific), myeloid immune populations (M1/M2 macrophages, monocytes, dendritic cells, mast cells, neutrophils), lymphoid populations (CD8+ cytotoxic T cells, exhausted T cells, CD4+ helper T cells, regulatory T cells, NK cells, B cells, plasma cells), and stromal populations (fibroblasts, cancer-associated fibroblasts, endothelial cells, pericytes). Tissue-specific epithelial signatures included luminal, basal, and HER2 programs for breast; alveolar, club cell, and ciliated programs for lung; colonocyte, goblet, stem cell, and Wnt pathway programs for colorectal; and astrocyte, oligodendrocyte, neural progenitor, and neuron programs for GBM. Each signature consisted of 3–8 established marker genes derived from published single-cell RNA-seq atlases: Neftel et al. [17] for GBM cell states, Wu et al. [18] for breast cancer, Travaglini et al. [19] for lung, and Lee et al. [20] for colorectal cancer. Immune and stromal markers were drawn from Newman et al. [8] and the Human Protein Atlas. Complete gene lists are provided in Supplementary Table S3.

### TME score computation

For each signature, the score was computed as the arithmetic mean of the log_2_-transformed expression values of its constituent marker genes present in the expression dataset. Gene coverage was assessed per cohort, with signatures requiring a minimum of 2 detected genes. Scores were z-score standardized within each cohort independently to account for platform-specific expression distributions. This within-cohort normalization is critical for cross-platform validation, as it ensures that the relative ordering of samples on each TME axis is preserved regardless of absolute expression scale.

### Machine learning models

Two classifiers were trained for each driver–cancer pair: (1) L2-regularized logistic regression (C=1.0, max iterations=2,000) and (2) gradient boosting classifier (100 estimators, max depth=3, minimum samples per leaf=10). Both models used TME signature z-scores as features (22–28 features per cancer type). Internal performance was estimated via 5-fold stratified cross-validation on the TCGA training cohort. The model with higher external validation AUC was selected for reporting. All models were implemented in scikit-learn (v1.3) with a fixed random seed of 42 for reproducibility.

### External validation procedure

For external validation, the full TCGA training cohort was used to fit the final model, which was then applied to predict driver mutation status in the external cohort. TME scores in the external cohort were computed and z-normalized independently (no information from the training cohort was used for normalization). Discriminative performance was assessed using the area under the receiver operating characteristic curve (AUC) with bootstrap 95% confidence intervals (1,000 iterations). Area under the precision-recall curve (AUPRC) and Brier scores were computed to assess performance under class imbalance and calibration quality, respectively. External AUC ≥0.65 was considered validated, 0.55–0.65 marginal, and <0.55 failed, consistent with thresholds used in prior mutation-prediction studies using expression or histology features. Calibration curves were generated using 8-bin calibration with observed versus predicted frequency plots.

### Sensitivity analyses

ssGSEA comparison. To assess robustness to scoring methodology, we re-computed TME scores using single-sample Gene Set Enrichment Analysis (ssGSEA) with an exponent α=0.25. ssGSEA uses a rank-based enrichment statistic that is independent of absolute expression scale. Cross-validation AUCs were compared between mean z-score and ssGSEA scoring across all driver– cancer pairs. Tumor purity correction. An ESTIMATE-based [13] purity proxy was computed as the negative sum of mean stromal marker expression (COL1A1, COL1A2, COL3A1, VIM, FN1, FAP, DCN, LUM) and mean immune marker expression (PTPRC, CD3D, CD3E, CD8A, CD4, MS4A1, CD19, CD68), z-normalized within each cohort. This score was added as an additional feature to the logistic regression model, and purity-corrected CV AUCs were compared to the original AUCs. Permutation testing. For each driver–cancer pair, significance was assessed using 200 permutations of the class labels with cross-validated AUC as the test statistic. The permutation p-value was computed as the fraction of permuted scores exceeding the observed score. ERBB2 circularity analysis. Because the HER2 program signature contains ERBB2 and co-amplified genes from the 17q12 amplicon, we retrained and re-validated the ERBB2 amplification model after removing the HER2 program feature to confirm that predictive performance was not driven by circular detection of the amplicon’s own gene expression. Negative control analysis. To confirm biological specificity, we trained 100 logistic regression models with randomly assigned binary labels (preserving the observed ERBB2 amplification prevalence) on the TCGA-BRCA training set, then evaluated each on the METABRIC external cohort against real ERBB2 amplification labels. The null distribution of AUCs was compared to the real model’s performance. Clinical utility metrics. For the best-performing model (ERBB2 amplification on METABRIC), we computed sensitivity, specificity, positive predictive value (PPV), and negative predictive value (NPV) at the Youden’s J optimal threshold and at a fixed threshold of 0.50.

### KRAS co-mutation subgroup analysis

KRAS-mutant LUAD tumors in TCGA were stratified by co-occurring STK11 and TP53 mutations into three subgroups: KRAS+STK11 (n=36), KRAS+TP53 (n=49, excluding STK11 co-mutants), and KRAS-only (n=69, neither STK11 nor TP53 co-mutant) [10]. KRAS wild-type tumors (n=348) served as the reference group. Mean TME z-scores were compared across subgroups, and pairwise significance was assessed using Mann–Whitney U tests.

### Survival analysis

Overall survival analysis was performed on METABRIC [9] using the lifelines Python package. The primary analysis used Cox proportional hazards regression with the continuous predicted probability as the sole covariate, providing a threshold-independent test of prognostic relevance. For visualization, patients were additionally stratified by predicted mutation probability at the median threshold, and Kaplan–Meier curves were compared using the log-rank test. Robustness to threshold choice was confirmed by repeating the stratified analysis at tertile cutpoints. Multivariate Cox proportional hazards models included TME-predicted probability as the primary covariate alongside age at diagnosis (continuous), histological grade (ordinal), and PAM50/Claudin molecular subtype (categorical, dummy-encoded with Basal as reference). Model discrimination was assessed by Harrell’s concordance index, comparing clinical-only versus clinical-plus-TME models.

### LM22 signature comparison

To evaluate the choice of custom tissue-specific signatures over established immune reference matrices, we scored TCGA-BRCA and METABRIC samples using the canonical LM22 immune cell signature matrix (22 immune cell types) [8]. For each BRCA driver, logistic regression models were trained separately on custom TME features, LM22 features, and combined features, and external validation AUC was compared across the three feature sets. Spearman correlation between custom and LM22 AUC rankings was computed to assess concordance.

### Feature attribution

SHAP (SHapley Additive exPlanations) [12] values were computed using LinearExplainer applied to the logistic regression models on external test sets. Feature reduction curves were generated by sequentially training models on the top-k features ranked by absolute logistic regression coefficient (k = 1, 2, 3, 5, 8, 10, 15, 20, all) and evaluating external validation AUC at each step.

### Multi-label prediction correlation

To assess whether independent driver prediction models captured overlapping or distinct biological signals, we computed pairwise Pearson correlations among predicted probabilities for all six BRCA drivers across 1,859 METABRIC samples.

### Statistical analysis of TME differences

For each validated driver in external cohorts, TME signature score differences between mutant and wild-type tumors were assessed using two-sided Mann–Whitney U tests. To account for multiple comparisons across signatures within each driver, p-values were corrected using the Benjamini–Hochberg procedure, and significance was defined at FDR q<0.05. Effect sizes were reported as mean z-score differences between groups. Nominal p-values and FDR-adjusted q-values are both reported in Supplementary Table S2.

### Software and reproducibility

All analyses were performed in Python 3.10 using pandas (v2.0), numpy (v1.24), scipy (v1.11), scikit-learn (v1.3), lifelines (v0.27), shap (v0.43), and matplotlib (v3.7). GEOparse (v2.0) was used for microarray platform annotations. Benjamini–Hochberg FDR correction was performed using scipy.stats.false_discovery_control. All code and signature definitions are available from the corresponding author upon reasonable request.

## Data Availability

The datasets analyzed during the current study are available in the cBioPortal for Cancer Genomics (https://www.cbioportal.org) and the NCBI Gene Expression Omnibus (https://www.ncbi.nlm.nih.gov/geo/) under accession numbers GSE72094 and GSE39582. CPTAC data are available via the Proteomic Data Commons. All analysis code is publicly available at https://github.com/earekab2006-hue/tme-genotyping.

## Acknowledgements

The authors received no specific funding for this work.

## Funding

This research received no specific grant from any funding agency in the public, commercial, or not-for-profit sectors.

## Author Contributions

E.B. conceived the study and designed the analysis framework. E.B. and N.M. developed the software pipeline. E.B. performed the computational analyses and survival modeling. E.B. wrote the manuscript. All authors reviewed and edited the manuscript.

## Competing Interests

The authors declare no competing interests.

## Supplementary Information

**Supplementary Table S1**. Complete internal cross-validation results for all driver–cancer pairs across four cancer types, including AUC, standard deviation, and permutation p-values.

**Supplementary Table S2**. TME signature differences between mutant and wild-type tumors in external validation cohorts.

**Supplementary Table S3**. Complete TME signature definitions including all marker genes for each cell-type program across four cancer types.

**Supplementary Table S4**. Mutation frequency comparison between TCGA training and external validation cohorts.

**Supplementary Table S5**. Multivariate Cox proportional hazards models for TME-predicted ERBB2 amplification and TP53 mutation scores in METABRIC.

**Supplementary Figure S1**. ERBB2 circularity analysis: comparison of full model versus model with HER2 program removed.

**Supplementary Figure S2**. Calibration curves for all 15 driver–cancer pairs on external validation cohorts.

**Supplementary Figure S3**. BRCA driver prediction correlation matrix showing pairwise Pearson correlations among predicted probabilities for six drivers across 1,859 METABRIC samples.

**Supplementary Figure S4**. KRAS co-mutation subgroup TME profiles in LUAD.

**Supplementary Figure S5**. SHAP feature attribution plots for the top four externally validated drivers.

**Supplementary Figure S6**. Comparison of external validation AUC using custom TME signatures versus LM22 immune-only signatures for all six BRCA drivers on METABRIC.

## Notes

https://github.com/earekab2006-hue/tme-genotyping

### Competing Interest Statement

The authors have declared no competing interest.

